# Addressing Artemisinin Resistance in Malaria Treatment: A Novel Ferrous ACT (FACT) Formulation

**DOI:** 10.1101/2024.02.15.579727

**Authors:** Yuliang Wang, Kexuan Tang

**Affiliations:** Frontiers Science Center for Transformative Molecules, Joint International Research Laboratory of Metabolic & Developmental Sciences, Plant Biotechnology Research Center, Fudan-SJTU-Nottingham Plant Biotechnology R&D Center, School of Agriculture and Biology, Shanghai Jiao Tong University, Shanghai 200240, China; Shanghai Jiao Tong University Sichuan Research Institute, Chengdu, 610213, China

## Abstract

Malaria is the worldwide infectious disease causing significant losses of human lives every year. The deployment of artemisinin-based combination therapies (ACTs) has helped to reduce the losses caused by malaria. However, resistance to ACTs evolved especially by *Plasmodium falciparum* occurs and becomes more and more threats for human lives. Up to now, there is no effective way to solve this drug-resistance problem. Here we propose a new ACT, which contains traditional ACT plus iron supplement, so-called ferrous ACT (FACT). The FACT is promising to become a safe and effective new formulation to combat ACT resistance malaria, helping WHO reach its 2030’s Malaria Control Goal.

## Background

In the era of artemisinin-based combination therapies (ACTs) emerging as the primary treatment for malaria, resistance has surfaced in clinical settings. Since 2008, there has been a rising prevalence of reduced sensitivity to artemisinin-type drugs among malaria parasites, notably in Southeast Asia [1]. Recent studies in 2021 have also identified the emergence of artemisinin-resistant malaria in Africa, which constitutes 90% of the global malaria burden [2]. The confirmed clinical resistance of malaria parasites to artemisinin has independently originated and spread locally in Africa. The development of resistance in malaria parasites to artemisinin is attributed to mutations in the Kelch13 gene [3]. The prevalence of malaria parasites carrying Kelch13 mutations has significantly increased, rising from 3.9% in 2015 to 19.8% in 2019. This escalation is primarily due to the increased frequency of allele genes such as A675V and C469Y [2]. Model predictions indicate that without intervention, the ineffectiveness of artemisinin and its derivatives in Africa could result in an additional 78 million cases and 116,000 deaths within five years.

These mutations result in a decrease in the activity of the Kelch13 protein and its interacting substances, thereby reducing the hemoglobin’s endocytosis process. Consequently, the activation of artemisinin and its derivatives (ART) is diminished, ultimately leading to resistance of the malaria parasite to artemisinin and its derivatives [4]. K13 gene mutations only cause a reduction in endocytosis during the ring stage of the malaria parasite, and when entering the trophozoite stage, it does not affect the endocytosis of hemoglobin. At this stage, sensitivity to artemisinin-type drugs is restored. To withstand artemisinin-type drugs, malaria parasites alter their lifecycle, lengthening the ring stage and shortening the trophozoite stage, leading to prolonged parasite clearance times. Conventional 3-day treatment regimens often result in treatment failure in regions with a high prevalence of K13 mutations. Professor Tu Youyou, a winner of the 2015 Nobel Prize in Physiology or Medicine, has proposed a temporizing solution to address the current artemisinin resistance epidemic by extending ACTs treatment to 7 days.

Previous studies have indicated that artemisinin-type drugs require activation by ferrous iron or heme *in vivo* [5]. Malaria parasites ingest hemoglobin into the digestive vacuole to obtain amino acids for growth and development, concurrently producing a significant amount of heme that activates artemisinin-type drugs. Malaria parasites reduce endocytosis through K13 gene mutations, decreasing heme production and artemisinin-type drug activation. The question arises whether this state can be altered by supplementing with iron.

## Methods

Artesunate, amodiaquine, ferrous sulfate, and heme iron raw materials were donated by Guilin Pharmaceutical Co., Ltd. (China). Eight-week-old male C57 mice were purchased from Shanghai Slac Laboratory Animal Co., Ltd. After one week of acclimatization, the mice were divided into four groups and orally administered by gavage: 1) Control group, administered with artesunate-amodiaquine compound, artesunate 1.67 mg/kg + amodiaquine 4.5 mg/kg; 2) Ferrous sulfate group, compound plus ferrous sulfate, artesunate 1.67 mg/kg + amodiaquine 4.5 mg/kg + ferrous sulfate 60 mg/kg; 3) Heme iron group, compound plus heme iron, artesunate 1.67 mg/kg + amodiaquine 4.5 mg/kg + heme iron 56 mg/kg; 4) Mixed iron group: compound plus ferrous sulfate and heme iron, artesunate 1.67 mg/kg + amodiaquine 4.5 mg/kg + ferrous sulfate 30 mg/kg + heme iron 28 mg/kg. The mice were continuously administered for 7 days, euthanized 2 hours after the last administration, and blood, liver, and kidney were collected. Blood levels of alanine aminotransferase (ALT) and aspartate aminotransferase (AST) were measured using an automatic biochemical analyzer. Additionally, concentrations of artesunate and dihydroartemisinin in both blood and liver tissues of mice were determined through high-performance liquid chromatography. Liver and kidney tissues were subsequently embedded in paraffin, sectioned, and stained with hematoxylin and eosin (HE) to assess cellular morphology. Notably, this experimental protocol received ethical approval from the Animal Ethics Committee of Shanghai Jiao Tong University.

## Results

Following seven consecutive days of administration, the combination of artesunate and amodiaquine compound therapy with ferrous sulfate or heme iron did not affect the levels of alanine aminotransferase (ALT) and aspartate aminotransferase (AST) in the blood of mice. Histopathological examination results also indicated that supplementation with iron agents did not induce additional oxidative stress reactions in the liver and kidneys of mice when using artemisinin-based combination therapy (ACT).

We utilized high-performance liquid chromatography to detect the levels of artesunate and its active metabolite dihydroartemisinin in the blood and liver tissues of mice 2 hours after drug administration. Given the short half-life of artemisinin compounds, artesunate or its metabolites were no longer detectable in the blood 2 hours after administration. The liver serves as the primary organ for drug accumulation and metabolism and is also the first parasitized organ following Plasmodium infection in the human body. Supplementation with different iron agents has varied effects on the accumulation and metabolism of ACT compounds in the liver. Ferrous sulfate significantly increased the levels of artesunate and its metabolite dihydroartemisinin in mouse liver tissue. Conversely, supplementation with heme iron reduced the levels of artesunate in mouse liver tissue, although it did not significantly affect the levels of its active metabolite, dihydroartemisinin.

## Discussion

The exact molecular mechanism of the impact of iron supplementation on malaria infection and treatment is not yet clear. Clinical observations suggest that iron-deficiency anemia can prevent severe malaria, and iron supplementation can increase susceptibility to clinical malaria. *In vitro* experiments also indicate that the efficiency of malaria parasites infecting iron-deficient red blood cells is low, and iron supplementation reverses the protective effect of iron deficiency on malaria infection [6]. Transferrin in young red blood cells reduces intracellular iron aggregation and the risk of malaria infection [7]. The latest review examined a plethora of clinical research findings and concluded that in malaria-endemic areas, timely and effective provision of malaria prevention and management services, including iron supplementation, does not increase the likelihood of clinical malaria infection or worsen malaria symptoms [8]. Populations in regions with a high prevalence of malaria often experience concurrent anemia. Following malaria infection, the consumption of red blood cells in the body further intensifies the severity of anemia, leading to more serious health issues. Therefore, adding iron supplements to artemisinin-based treatments, forming a new combination formulation (Ferrous ACT, FACT) consisting of artemisinin-type drugs, other antimalarial drugs, and iron supplements, can address both anemia associated with malaria and potentially overcome the current resistance issues through ferroptosis mechanisms. Iron is a vital and essential element in life, influencing various physiological and biochemical processes. Recent research has focused on the relationship between ferroptosis and various diseases. Ferroptosis involves the production of lipid peroxidation and elevated reactive oxygen species (ROS) through the Fenton reaction in cells, leading to programmed cell death. Studies have shown that host liver cells can control Plasmodium liver-stage infection by regulating the SLC7a11-GPX4 signaling pathway to induce lipid peroxidation and ROS [9]. Low-dose dihydroartemisinin combined with ferroptosis inducers synergistically enhances antimalarial effects, while ferroptosis inhibitors had an obvious antagonistic effect [10]. Intracellular iron concentration is a complex balance, and sufficient iron supply enhances Plasmodium clearance through ferroptosis pathway and restoration of immune cell function. However, this may also lead to excessive immune damage [11]. Based on the typical usage of ACTs and iron supplements, we determined the daily dosage of artemisinin-based combinations with different iron supplements for mice. After continuous administration for 7 days, we observed the safety of the FACT formulation in mice, revealing no significant toxic reactions in the liver and kidneys (Figure 1). Therefore, FACT is promising to become a safe new formulation to combat artemisinin resistance.

**Figure 1:**
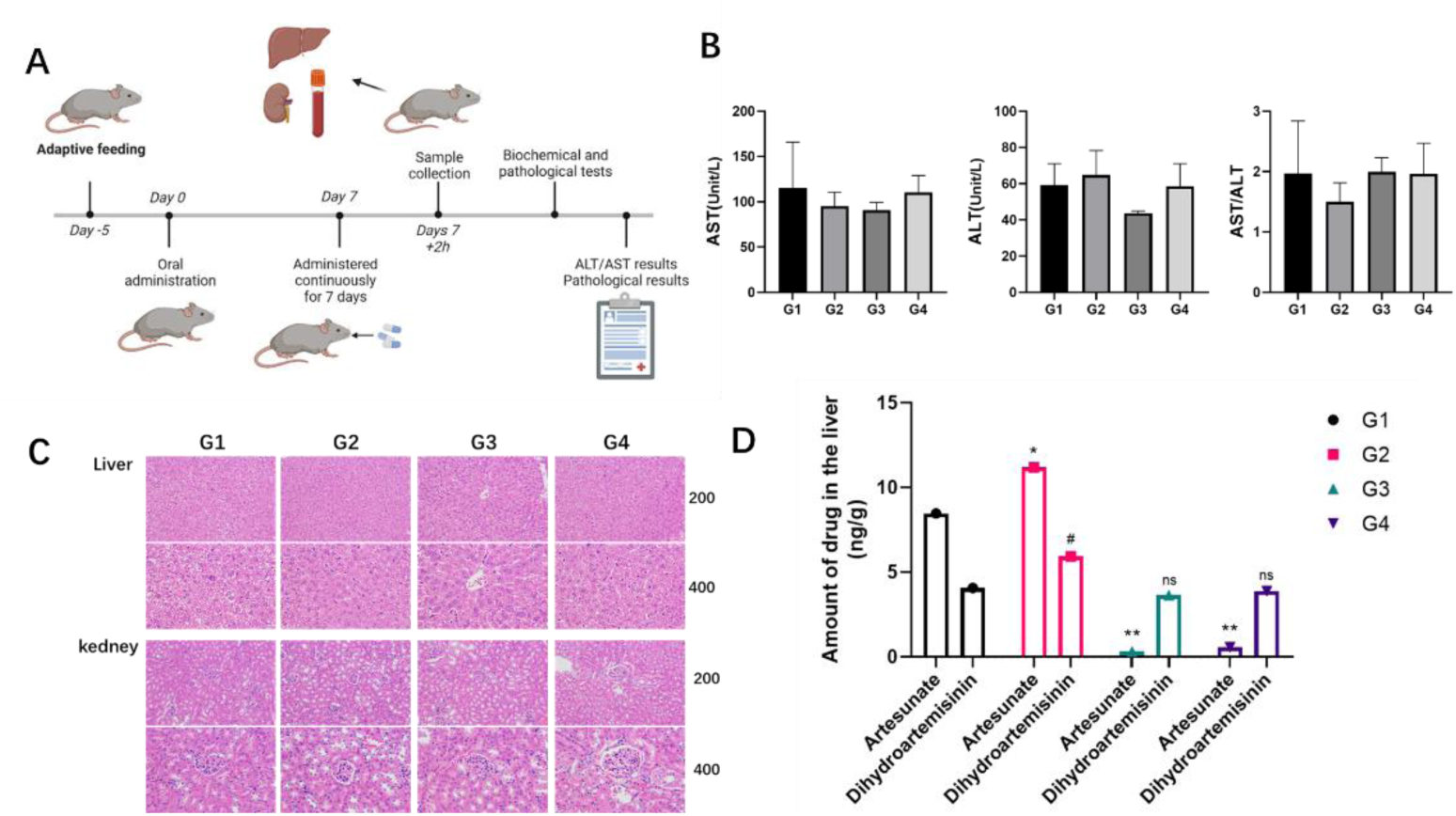
Safety results of mice orally administered artesunate-amodiaquine in combination with different iron supplements for 7 consecutive days. G1: artesunate 1.67 mg/kg + amodiaquine 4.5 mg/kg; G2: artesunate 1.67 mg/kg + amodiaquine 4.5 mg/kg + ferrous sulfate 60 mg/kg; G3: artesunate 1.67 mg/kg + amodiaquine 4.5 mg/kg + hematinic Iron 56 mg/kg; G4: artesunate 1.67 mg/kg + amodiaquine 4.5 mg/kg + ferrous sulfate 30 mg/kg + hematinic Iron 28 mg/kg **A**. Timeline of drug administration and sampling in mice. **B**. Levels of ALT and AST in mouse blood. The addition of iron supplements to artesunate-amodiaquine had no significant impact on ALT and AST levels. **C**. Histological results of HE staining in mouse liver and kidney sections. The addition of iron supplements to artesunate-amodiaquine showed no apparent effects on the liver and kidney tissues of mice. D. The content of artesunate and dihydroartemisinin in mouse liver tissue. Artesunate (ART), Dihydroartemisinin (DHA).

According to the World Health Organization (WHO)’s 2030 Global Malaria Technical Strategy and Goals, malaria treatment is a crucial component of this plan. Whether Professor Tu Youyou’s proposal to extend the ACT treatment period, the successful group dosing strategy in some countries, or our proposed FACT solution to address artemisinin resistance, all require an ample and affordable supply of artemisinin raw materials. Based on synthetic biology strategies, we have developed new varieties of artemisinin with a content as high as 3.2%, more than double that of existing conventional varieties [12]. With the support of the Bill and Melinda Gates Foundation, we are collaborating with scientists worldwide to achieve yeast-based total synthesis of artemisinin. These efforts offer potential solutions to the accessibility of artemisinin raw materials. We believe that the entire artemisinin industry chain, from research, cultivation, extraction to the production of combination formulations, all share the common goal of eradicating malaria, and we will eventually realize the global dream of a malaria-free world.

## Acknowledgements

This work was supported, in whole or in part, by the Bill & Melinda Gates Foundation (grant number: INV-027291). Under the grant conditions of the Foundation, a Creative Commons Attribution 4.0 Generic License has already been assigned to the Author Accepted Manuscript version that might arise from this submission.

